# Defining the mechanisms of action and resistance to the anti-PD-1+LAG-3 and anti-PD-1+CTLA-4 combinations in melanoma flank and brain models

**DOI:** 10.1101/2023.04.14.536907

**Authors:** Manali S. Phadke, Jiannong Li, Zhihua Chen, Paulo C. Rodriguez, J.K. Mandula, Lilit Karapetyan, Peter A. Forsyth, Y. Ann Chen, Keiran S.M. Smalley

## Abstract

**Background:** Although the anti-PD-1+LAG-3 and the anti-PD-1+CTLA-4 combinations are effective in advanced melanoma it remains unclear whether their mechanisms of action and resistance overlap.

**Methods:** We used single cell (sc) RNA-seq, flow cytometry and IHC analysis of responding SM1 and B16 melanoma flank tumors and SM1 brain metastases to explore the mechanism of action of the anti-PD-1+LAG-3 and the anti-PD-1+CTLA-4 combination. CD4+ and CD8+ T cell depletion and ELISPOT assays were used to demonstrate the unique role of CD4+ T cell help in the anti-tumor effects of the anti-PD-1+LAG-3 combination. Tetramer assays confirmed the loss of CD8+ tumor-reactive T cells in brain tumors resistant to the anti-PD-1+LAG-3 combination.

**Results:** The anti-PD-1+CTLA-4 combination was associated with the infiltration of FOXP3+ regulatory CD4+ cells (Tregs), fewer activated CD4+ T cells and the accumulation of a subset of IFNγ secreting cytotoxic CD8+ T cells, whereas the anti-PD-1+LAG-3 combination led to the accumulation of CD4+ T helper cells that expressed CXCR4, TNFSF8, IL21R and a subset of CD8+ T cells with reduced expression of cytotoxic markers. T cell depletion studies showed a requirement for CD4+ T cells for the anti-PD-1+LAG-3 combination, but not the PD-1-CTLA-4 combination at both flank and brain tumor sites. In anti-PD-1+LAG-3 treated tumors, CD4+ T cell depletion was associated with fewer activated (CD69+) CD8+ T cells, impaired IFNγ release and increased numbers of myeloid-derived suppressor cells (MDSCs) but, conversely, increased numbers of activated CD8+ T cells and IFNγ release in anti-PD-1+CTLA-4 treated tumors. Analysis of relapsing melanoma brain metastases from anti-PD-1+LAG-3 treated mice showed an increased accumulation of MDSCs and a loss of gp100+ tumor reactive CD8+ T cells. An analysis of the inferred cell-cell interactions from the scRNA-seq data suggested the MDSCs interacted with multiple subsets of T cells in a bi-directional manner.

**Conclusions:** Together these studies suggest that these two clinically relevant ICI combinations have differential effects upon CD4+ T cell polarization, which in turn, impacted cytotoxic CD8+ T cell function. Further insights into the mechanisms of action/resistance of these clinically-relevant ICI combinations will allow therapy to be further personalized.

## Introduction

The development of immune checkpoint inhibitors (ICIs) has revolutionized the treatment of advanced melanoma (1). At this time, the most widely used ICIs are antibodies that block the function of programmed cell death protein (PD)-1, reversing the exhaustion of CD8+ cytotoxic T cells (2). These therapies, including nivolumab and pembrolizumab, are widely used for the treatment of melanoma and although successful, are frequently associated with both intrinsic and acquired resistance (3, 4). Two of the most widely explored additional ICIs are antibodies that target cytotoxic lymphocyte associated protein (CTLA)-4 and lymphocyte-activation gene (LAG)-3, both of which are upregulated upon T cell activation and associated with eventual T cell exhaustion. Inhibition of CTLA-4 is associated with low response rates in the single agent setting (5), with improved rates of response reported when CTLA-4 targeted therapies are used in combination with anti-PD-1 (6, 7), albeit at the cost of increased immune toxicity.

LAG-3 is a homolog of CD4 that binds to major histocompatibility complex (MHC) Class 2 (8). Preclinical studies demonstrated that co-targeting of LAG-3 and PD-1 frequently led to the regression of established tumors that were otherwise resistant to ICI monotherapy (9). In a recent randomized phase 2/3 clinical trial, the anti-PD-1+LAG-3 combination (nivolumab + relatlimab) significantly improved progression-free survival (PFS) compared to single agent nivolumab (10). Significantly, the combination was well tolerated with far fewer adverse events reported than in patients treated with the anti-PD-1+CTLA-4 combination. The anti-PD-1+LAG-3 combination was subsequently explored in the neoadjuvant setting for patients with resectable stage III or stage IV oligometastatic melanoma where the combination therapy was associated with pathological complete response rate (pCR) of 57% (11). In light of these encouraging findings, the anti-PD-1+LAG-3 combination was FDA approved for advanced melanoma in the Spring of 2022.

The brain is one of the most common sites of melanoma metastasis, with 40-60% of patients with advanced melanoma showing evidence of central nervous system (CNS) involvement (12, 13). Although it was previously believed that the CNS exists as an immune-privileged site, it is now understood that the brain and the spinal cord are under continual immune surveillance. However, the mechanisms by which pro- and anti-tumor immune subsets traffick into the CNS and the impact these subsets have on CNS-metastases remains incompletely understood. A recent report demonstrated that melanoma brain metastases (MBMs) had a lower level of T cell infiltration and micro-vessel density than metastases from extracranial sites (14). A second histological study also confirmed the lower level of CD8+ T cell infiltrate in MBM compared to matched extracranial metastases (15). Although the mechanisms underlying the reduced infiltration of T cells in MBM are not clear, there is some evidence from neuroinflammation studies that have suggested activated microglia may limit T-cell accumulation (16). In mouse MBM models, immunotherapy responses in the brain are dependent upon the peripheral expansion of T cells that traffic into the brain (17). Studies on matched cranial and extracranial melanoma metastases have demonstrated reduced T cell clonality in the brain compared to extracranial metastatic sites (18). We recently used single cell (sc) RNA-Seq analyses of MBM and melanoma skin metastases to demonstrate that the extent of T cell infiltration into the brain was lower than of the skin metastases (19). Other work demonstrated that MBM had a higher proportion of monocyte-derived macrophages and TOX+ CD8+ T cells with unique patterns of immune checkpoint expression compared to extracranial metastases (20).

Recent studies from our group have shown that >50% of MBM patients exhibit significant clinical intracranial responses to the combination of the immune checkpoint inhibitors (ICI) that target CTLA-4 and PD-1 (ipilimumab + nivolumab) (21). However, almost half of the treated population do not respond to treatment and the response rate and survival in patients with symptomatic MBM is significantly lower (∼20%) (21). The combination of relatlimab with nivolumab is currently being explored clinically in patients with MBM (NCT05704647). As little is known about its activity against MBM, or its mechanism of action in the brain, we here used dual flank/brain injection models of MBM to determine the mechanism of action of the anti-PD-1+LAG-3 and anti-PD-1+CTLA-4 combinations. Our studies demonstrated differential effects of the two combinations on CD8+ cytotoxic and CD4+ T cell populations, and the dependency of the anti-PD-1+LAG-3 combination upon a CD4+ T helper phenotype. Failure of anti-PD-1+LAG-3 therapy in the brain was associated with increased accumulation of myeloid derived suppressor cells (MDSCs) and a loss of tumor-reactive T cells.

## Material and Methods

### Cell lines and mice

SM1 mouse BRAF-mutant melanoma cells were obtained from Dr. Eric Lau, Moffitt Cancer Center. B16 mouse melanoma cells were obtained from Dr. Mary Jo Turk, Dartmouth University. The cells were maintained for maximum 10 passages in RPMI1640 + 10% FBS for SM1 and RPMI1640 + 7.5% FBS for B16 cells. Cell lines were routinely tested for *Mycoplasma* (every three months). Female immunocompetent C57BL/6J (The Jackson Laboratory) mice were observed daily, and all the protocols were reviewed and approved by Institutional Animal Care and Use Committee at University of South Florida (approved IACUC # 8889R).

### *In vivo* procedures

Female 7-week-old immunocompetent C57BL/6J mice were used for the experiments. All the protocols were reviewed and approved by Institutional Animal Care and Use Committee at University of South Florida (approval #8889R). Mice were subcutaneously injected with 1.5 × 10^6^ SM1 or B16 cells in Matrigel (cat. #CB40234; Fisher Scientific) into the right flank. Next day, these mice were injected with 50,000 SM1 cells in PBS into the caudate nucleus of the right cerebral hemisphere of the brain using stereotactic surgical procedures. The flank tumors were allowed to grow approximately to ∼ 50-70 mm^3^ and brain tumors were allowed to grow until visible via MRI, before initiation of drug dosing. For the B16 model, the drugs dosing was initiated next day after the injections. Mice received intraperitoneal doses of anti–PD-1 antibody (200 μg/100 μL; clone RMP1-14; cat. #BE0146; Bio X Cell) or anti-CTLA-4 (200 μg/100 μL; clone 9D9; cat. #BE0164; Bio X Cell) or anti-LAG-3 (200 μg/100 μL; clone C9B7W; cat. #BE0174; Bio X Cell) as a single agent or as a combination every 5 days. For isotype control, a cocktail of IgG2a isotype control (200 μg/100 μL; clone 2A3; cat. #BE0089; Bio X Cell), IgG1 isotype control (200 μg/100 μL; clone HRPN; cat. #BE0088; Bio X Cell) and Syrian hamster IgG (200 μg/100 μL; cat. #BE0087; Bio X Cell) were administered intra-peritoneally every 5 days. For the B16 model, mice received the antibodies every day. The flank tumors were measured using calipers and the brain tumors were monitored and measured by MRI scanning. The tumors were collected at the endpoint, weighed, and processed for single-cell RNA sequencing (scRNA-seq), flow cytometry analysis or immunohistochemistry.

### CD4^+^ and CD8^+^ T cell depletion

CD4-specific antibody (clone YTS191; cat. #BE0003-1; Bio X Cell) and CD8a-specific antibody (clone YTS169.4; cat. #BE0117; Bio X Cell) from Bio X Cell were used to deplete CD4^+^ T cells and CD8^+^ T cells, respectively. The C57BL/6J mice were administered anti-CD4 and anti-CD8a (100 μg/100 μL) via intraperitoneal injections three days before injection with SM1 or B16 cells and then every 4 days thereafter. When the flank tumors reached 50-70 mm^3^ in size and brain tumors were visible by MRI, the mice were treated with either IgG control, anti-PD1+ LAG-3 or anti-PD1+CTLA-4 every 5 days. Tumor size was measured twice weekly, with CD4^+^ T-cell or CD8^+^ T-cell depletion measured by flow cytometry at termination of the experiment.

### Flow cytometry

Tumors were harvested at the endpoint or at listed time points for the kinetics experiments, under sterile conditions and weighed. Single-cell suspensions were prepared by enzymatic digestion, using a MACS tumor dissociation kit (cat. #130-095-929; Miltenyi Biotec) and the number of viable cells counted. To analyze immune-cell populations, 1 × 10^6^ cells were blocked with purified mouse CD16/32 antibody (1:100 dilution; cat. #101301; BioLegend) for 5 minutes on ice. The cells were then incubated with antibody cocktail of Live/Dead Near IR antibody (cat. #L10119; Thermo Fisher Scientific), anti-CD45-BUV395 (clone 30-F11; cat. #565967; BD Biosciences), anti-CD3-BUV737 (clone 17A2, cat. #564380; BD Biosciences), anti-CD4-BUV 496 (clone GK1.5; cat. #564667; BD Biosciences), anti-CD8-BUV805 (clone 53-6.7; cat. #564920; BD Biosciences), CD127-BV711 (clone SB/199; cat. #565490; BD Biosciences), CD69-AF488 (clone H1.2F3; cat. # 104516; BioLegend), CD44-APCR700 (clone IM7; cat. #565480; BD Biosciences), CD62L-BV650 (clone MEL-14; cat. #564108; BD Biosciences), PD-1-BV785 (clone 29F.1A12; cat. #135225; BioLegend), CTLA-4-BV421 (clone UC10-4B9; cat. #106311; BioLegend), TIM3-PECF594 (clone B8.2C12; cat. #134013; BioLegend), LAG-3-PE (clone C9B7W; cat. #552380; BD Biosciences) for T-cell analysis and Live/Dead Near IR antibody (cat. #L10119; Thermo Fisher Scientific), anti-CD45-BUV395 (clone 30-F11; cat. #565967; BD Biosciences), anti-CD3-BUV737 (clone 17A2, cat. #564380; BD Biosciences), CD11b-BB700 (M1/70, cat. #566417; BD Biosciences), Gr-1-PE-Cy7 (RB6-8C5; cat. #108415; BioLegend), anti-Ly6C-BV421 (clone HK1.4; cat. #128031; BioLegend), anti-Ly6G-APC (clone 1A8; cat. #127613; BioLegend), anti-CD11c-BV605 (clone N418; cat. #117333; BioLegend), anti-MHC II-BB515 (clone 2G9; cat. #565254; BD Biosciences), anti-F4/80-BV785 (clone BM8; cat. #123141; BioLegend), and anti-CD103-PE (clone M290; cat. #561043; BD Biosciences) for myeloid cells analysis. Antibodies were used according to the manufacturer’s instructions. The cells were incubated with the antibody cocktail for 20 minutes at 4°C in dark. For FOXP3 staining, the cells were fixed/permeabilized overnight at 4°C in dark using eBioscience FOXP3 transcription factor staining buffer set (cat. #00-5523-00; Thermo Fisher) and FOXP3 monoclonal antibody (1:25 dilution; clone FJK-16s; cat. #17-5773-80; Thermo Fisher). All the washings were done with PBS (cat. #SH30256FS; Fisher Scientific) + 2% FBS (cat. #F0926; Sigma). Flow cytometry acquisition was performed on the BD FACS Symphony or LSR II. The data analysis was carried out using FlowJo software. To detect melanoma antigen expression on tumor cells or cell lines treated with kinase inhibitors, the cells were incubated with anti-Tyrp1-APC (clone TA99; cat. # NBP2-34720APC; Novus Biologicals) at 1:50 dilution for 20 minutes at 4°C in dark. Flow cytometry acquisition was performed on the BD FACS LSR II. The data analysis was carried out using FlowJo software.

### gp-100 Tetramer assay

The number of gp-100 TCR positive cells in MBM samples from mice responding and relapsing on anti-PD1+LAG-3 and anti-PD1+CTLA-4 therapy was determined by carrying out *in vitro* gp-100 tetramer assays. Prior to staining by gp-100 tetramer, a single cell suspension of cells from MBMs of responding and relapsing mice was prepared by enzymatic digestion, using a MACS tumor dissociation kit (cat. #130-095-929; Miltenyi Biotec). The number of viable cells were counted, and around 1×10^6^ cells were blocked with purified mouse CD16/32 antibody (1:100 dilution; cat. #101301; BioLegend) for 5 minutes on ice. The cells were then stained with gp-100 tetramer PE (MBL #TS-M505-1 H-2D^b^ human gp100 tetramer KVPRNQDWL-PE) and incubated for 30 mins. Additionally, the cells were also stained with antibody cocktail of Live/Dead Near IR antibody (cat. #L10119; Thermo Fisher Scientific), anti-CD45-BUV395 (clone 30-F11; cat. #565967; BD Biosciences), anti-CD8-BUV805 (clone 53-6.7; cat. #564920; BD Biosciences). Flow cytometry acquisition was performed on the BD FACS LSR II. The data analysis was carried out using FlowJo software.

### IFNγ ELISPOT assay

IFNγ ELISPOT assay was performed using Mouse IFN-gamma ELISpot kit (cat. # EL485; R&D systems) according to the manufacturer’s instructions. Around 1 million cells from SM1 and B16 tumors were stimulated with eBioscience Cell Stimulation Cocktail (500X) (cat. #00-4970-03; Thermo Fischer Scientific) and plated in the 96-well plate provided overnight at 37°C in a 5% CO2 incubator. The assay was performed as per the manufacturer’s instructions.

### sc-RNA seq

Tumors were harvested at the endpoint under sterile conditions and weighed. Single-cell suspensions were prepared by enzymatic digestion, using a MACS tumor dissociation kit (cat. #130-095-929; Miltenyi Biotec). Cells were strained through MACS strainer (cat. # 130-098-458; Miltenyi Biotec). The cell count and viability were analyzed by staining the cells with AO/PI stain on the Nexcelom Cellometer K2. The cells were then resuspended at a concentration of 500 cells/μL in PBS (cat. #SH30256FS; Fisher Scientific) + 0.4% nonacetylated BSA (cat. #BP1605100; Fisher Scientific). The samples were then loaded onto 10X Genomics Chromium Single-Cell Controller (10X Genomics) to prepare scRNA-seq libraries. Around 50,000 to 1,000,000 mean sequencing reads per cell were generated on Illumina NextSeq 500 instrument using v2.5 flow cells. 10X Genomics CellRanger software was used for demultiplexing, barcode processing, alignment, and gene counting. Finally, the analysis of single-cell data set was performed using Interactive Single-Cell Visual Analytics (ISCVA) which was previously outlined in (22, 23). Cells with high mitochondria content were not filtered as these may reflect cell populations going through apoptosis. Data are available through Gene Expression Omnibus (GEO: pending).

### Cell-cell interaction analysis

To investigate cell–cell interaction (CCI) mediated by ligand-receptor complexes, especially between T cells and MDSCs, SingleCellSignalR was used (24). SingleCellSignalR predicts ligand (L) and receptor (R) interactions between two cell types using a regularized LRscore based on their curated LR database which contains 3,251 L–R interactions. LRscore is a scaling product score of average ligand expression in one cell type and average receptor expression in the other cell type. In pairs of cells with statistically significant differences using the nominal *P* value of 0.05 as threshold, we further investigated each specific interaction using Wilcoxon test.

### sc-VDJ sequencing and clonal diversity analysis

TCR reads sequenced by V(D)J assay were aligned to mouse reference transcriptome using Cell Ranger VDJ (v6.1, 10X Genomics). T cells were assigned with productive CDR3 regions for TRA/TRB chains. Each clonotype was assigned with a unique identifier based on amino acid sequences of the CDR3 regions and V(D)J genes of the chains. To evaluate the clonality of TCR repertoire and clonal diversity, Chao 1 was calculated using the functions repDiversity in the R package “immunarch”13 (version 0.9.0). Chao1 is an tool for estimating the species richness asymptotically (25).

### Statistical analysis

One-way ANOVA in Microsoft Excel version 15.40 was used to compare the results between different groups with a single independent variable. The mean of three independent experiments SEM is shown for each data set. Results with P values ≤ 0.05 were considered statistically significant.

## Results

### Anti-PD-1+LAG-3 and anti-PD-1+CTLA-4 combinations are both effective against mouse models of melanoma flank tumors and MBM

As the relative effectiveness of the anti-PD-1+LAG-3 and anti-PD-1+CTLA-4 combinations have never been directly compared in flank and MBM models, we undertook studies in which mice were injected with SM1 melanoma cells carrying a BRAF mutation (BRAF^V600E^) and deletion in CDKN2A into the flank and stereotactically into the brain. A dual injection model was used as our preliminary studies had indicated that mice with brain metastases had poor immune responses and showed little response to ICI therapy (data not shown). Animals were treated with either IgG control, anti-PD-1, anti-LAG-3, anti-CTLA-4, anti-PD-1+CTLA-4 or anti-PD-1+LAG-3 (**Figure 1A**). Combination ICI therapies were more effective than each single agent ICI; notably 100% of flank tumors exhibited full regression in response to the anti-PD-1+LAG-3 and anti-PD-1+CTLA-4 combination therapy. Additionally, after flank tumor regression tumors did not regrow during the course of the experiment. Although the combination ICI therapies were also effective against brain metastases, 2/10 mice from each combination group exhibited tumor progression in the brain (**Figure 1B**). In mice exhibiting tumor regression in response to combination ICI therapy, tumor regrowth was not observed despite cessation of therapy; collectively, these findings indicate that combination ICI treatment can provoke curative, durable effects (**Figure 1C**). Next mice exhibiting tumor regression were rechallenged with an additional SM1 flank tumor. After 63 days SM1 rechallenged mice did not exhibit establishment of new tumors, indicating development of an anti-tumor immunological memory response (**Figure 1D,E**). We next evaluated the two ICI combinations in a more resistant B16 melanoma flank model and noted that although both combinations were initially effective at reducing the volume of established tumors (**Figure 1F,G,H**), 3/10 tumors relapsed on anti-PD-1+LAG-3 therapy. The anti-PD-1+CTLA-4 combination was also not curative in every mouse with 1/10 initially failing and 2/10 mice relapsing after cessation of treatment (**Figure 1H**). 7/10 of mice treated with anti-PD-1+CTLA-4 and 6/10 tumors on anti-PD-1+LAG-3 therapy did not relapse (**Figure 1H**). Mice responding to each of the ICI combinations were later rechallenged with tumor at day 97, with no regrowth detected (**Figure 1H**). It therefore seemed that although the B16 model exhibited greater resistance to combination ICI therapy, some mice did have prolonged anti-tumor responses to both anti-PD-1+LAG-3 and anti-PD-1+CTLA-4, associated with long term immunological memory.

**Figure 1:**
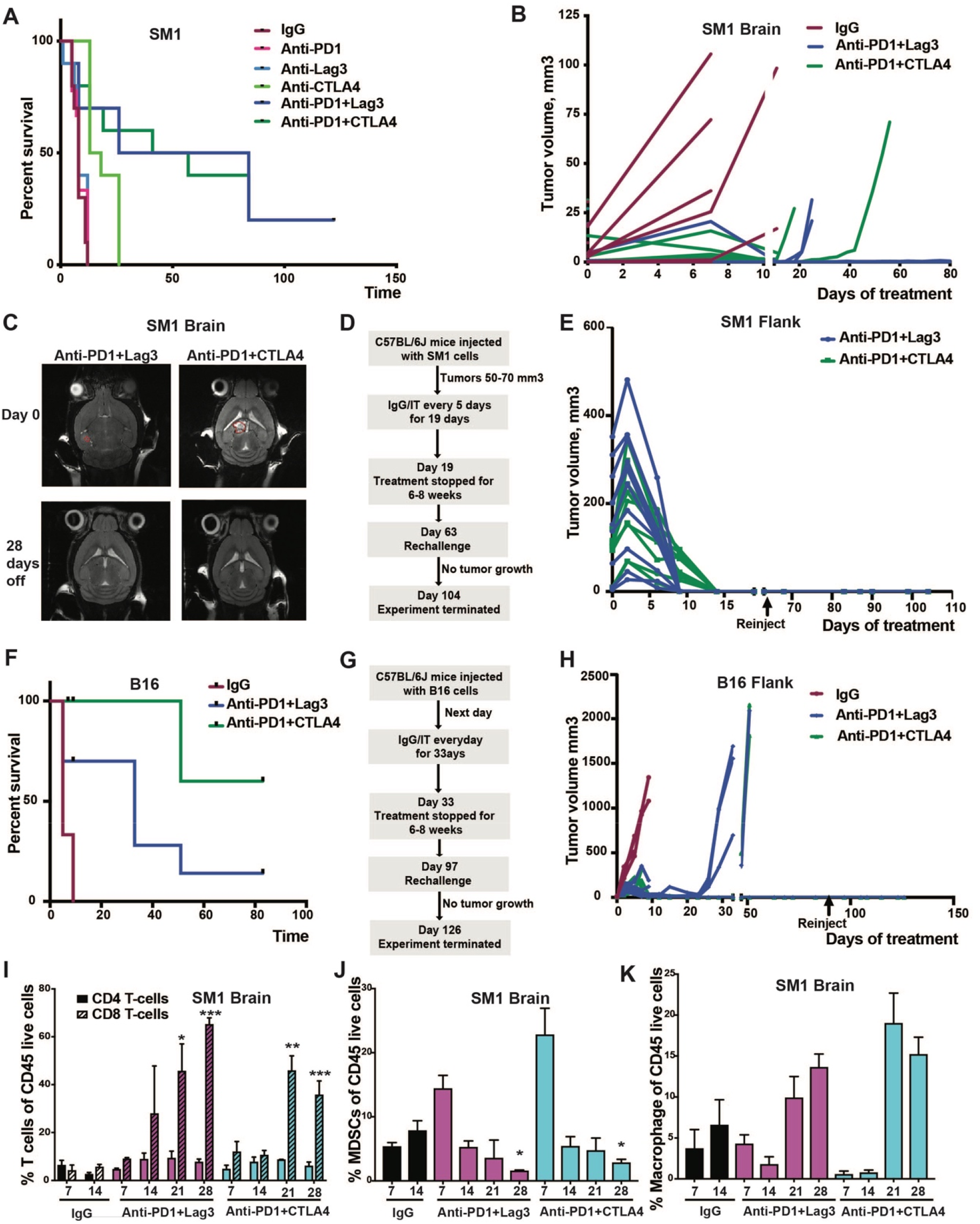
Anti-PD-1+LAG-3 and Anti-PD-1+CTLA-4 combinations suppress melanoma flank tumors and brain metastases in mouse melanoma models. A) Survival curves in SM1 mouse model for single agents Anti-PD1, Anti-LAG-3, Anti-CTLA-4 and the Anti-PD1+LAG-3 and Anti-PD1+CTLA-4 doublet combinations. B) The doublets were more effective than single agent ICI in the brain tumors of SM1 mouse model with 2/10 mice relapsed in each combination group. Mice received 200 μg/100 μL i.p. doses of single agents and doublets every 5 days for 80 days. C) MRI brain scans of SM1 mice demonstrating no tumor regrowth after holding the therapy for 30 days. D) Dosing schema for rechallenge experiment in SM1 mouse model. E) The ICI doublets were more effective than single agents ICI in the flank tumors of SM1 mouse model with no tumor regrowth during the experiment. Mice received 200 μg/100 μL i.p. doses of single agents and doublets every 5 days. F) Survival curves in B16 melanoma model for the anti-PD1+LAG-3 and Anti-PD1+CTLA-4 combinations. G) Dosing schema for rechallenge experiment in B16 melanoma mouse model. H) The doublets showed initial response in the flank tumors of B16 melanoma model with 3/10 tumors relapsed on anti-PD1+LAG-3 and 2/10 tumors relapsed on Anti-PD1+CTLA-4 after cessation of treatment. Mice received 200 μg/100 μL i.p. doses of combinations every day. I, J, K) Kinetics of CD4+ and CD8+ T cell infiltrate following treatment with each ICI doublet over a period of 28 days in SM1 brain tumors. The results were represented as average ± SEM with 10 mice per group for all the growth curve experiments. The results were represented as average ± SEM of 3 mice per group for panels I, J, K. Statistical significance was assessed with one-way ANOVA test (*, 0.001 ≤ p ≤ 0.05, **, 0.0001 ≤ p ≤ 0.001 and ***, p ≤ 0.0001).

To explore the kinetics of T cell accumulation in MBM following anti-PD-1+LAG-3 or anti-PD-1+CTLA-4 therapy we collected samples of responding SM1 brain tumors over time (day 7, 14, 21 and 28 days) (**Figure 1I,J,K**) and observed that levels of CD4+ and CD8+ T cell infiltration increased following treatment with each of the combinations. Of note, T cell infiltration was more rapid in response to anti-PD-1+LAG-3 (>14 days) whereas slower dynamics (>21 days) were noted to the anti-PD-1+CTLA-4 combination (**Figure 1J**). In the brain, initial levels of MDSC-like cells were high and then declined in response to both ICI combinations, whereas levels of macrophages increased (**Figure 1K**). We also collected samples from responding flank tumors over multiple time points (7 and 14 days) (**Supplemental Figure 1**). A limited range of time points was included for these analyses as the tumors regressed fully by day 15, leaving little tumor to analyze. It was noted that both the combination of anti-PD-1+LAG-3 and anti-PD-1+CTLA-4 led to increased infiltration of CD4+ and CD8+ T cells between day 7 and 14. In contrast, the IgG treated controls showed initially higher levels of CD4+ and CD8+ T cells that declined by day 14. An analysis of myeloid cells demonstrated the anti-PD-1+LAG-3 combination to increase accumulation of MDSC-like cells (**Supplemental Figure 1**) to a much greater extent than anti-PD-1+CTLA-4 combination or IgG control. Tumors from IgG treated mice saw increased numbers of macrophages over time, whereas treatment of melanomas with either anti-PD-1+CTLA-4 or anti-PD-1+LAG-3 led to a decrease in macrophage numbers.

### Single cell analysis of the immune landscape of flank and brain tumors treated with different combination immunotherapies

We next used droplet-based scRNA-seq to interrogate the immune landscape of SM1 tumors treated with either single agent or combination ICI. Analysis of samples from both responding flank tumors and resistant brain metastases (collected at day 25 for anti-PD-1+LAG-3 and day 19 for anti-PD-1+CTLA-4 treated tumors) along with detailed cell curation revealed a diverse cellular landscape composed of tumor cells, stromal cells (including fibroblasts, endothelial cells, micoglial cells) and multiple immune cell populations (T cells, NK cells, B cells, dendritic cells, monocytes, granulocytes, macrophages/MDSCs) (**Figure 2A**). The immune landscape of tumors treated with the PD-1+LAG-3 combination had a large proportion of T cells, the greatest accumulation of B cells, granulocytes and endothelial cells and the lowest proportion of macrophages. The anti-PD-1+CTLA-4 combination was also associated with an increased proportion of T cells, and the highest number of regulatory T cells (Tregs) (**Figure 2B**).

**Figure 2:**
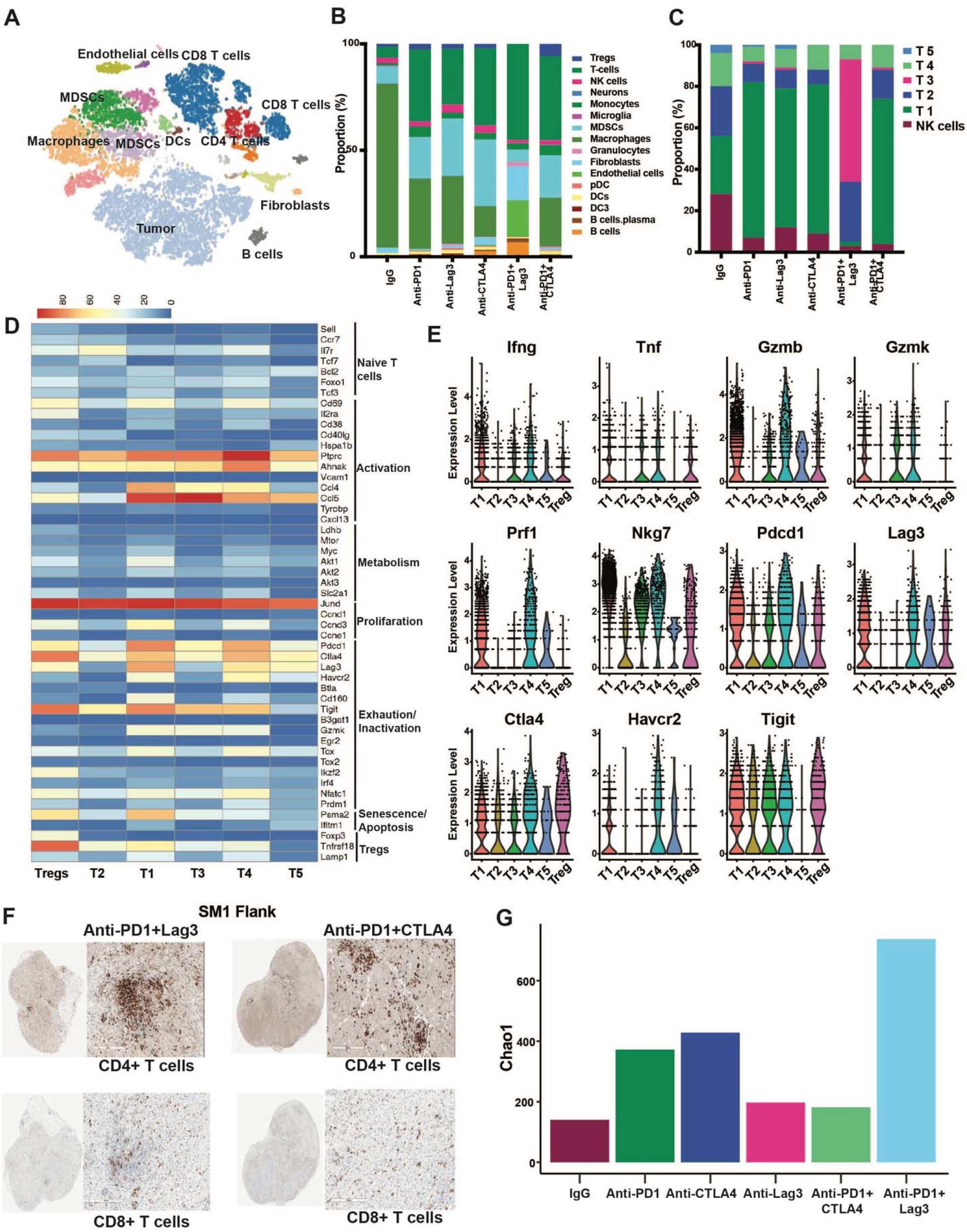
Immune landscape of flank and brain tumors in SM1 mouse model following the treatment with single agent and combination ICI therapy. A) t-SNE plots showing major cell types identified in relapsed brain tumors and responding flank SM1 tumors following treatment with single agent and combination ICI. B) Proportion of each cell type in the responding flank tumors from the indicted treatment groups. C) Proportion of different T-cell clusters identified in the responding flank tumors from the indicated treatment groups. D) Heatmap showing expression of activation/exhaustion markers across the identified T-cell clusters. E) Violin plots showing expression of T cell activation markers and immune checkpoints in each T-cell cluster. F) IHC analysis of tumors treated with anti-PD1+LAG-3 or PD-1+CTAL-4 demonstrates increased CD4+ and CD8+ T cell infiltration in responding flank tumors. The samples were stained with anti-CD4 and anti-CD8 by IHC. G) Chao1 analysis from the indicated treatment groups indicates the a major expansion of limited numbers of T cell clones in PD-1+LAG-3 treated SM1 tumors.

We next defined the transcriptional makeup of the T cells in tumors responding to single or combination immunotherapy (**Figure 2C**) and identified 6 major clusters including 4 sub-groups of CD8+ T cells (T1, 3, 4 and 5), one major subgroup of CD4+ T cells (T2) and NK cells. It was noted that the anti-PD-1+CTLA-4 combination was associated with the largest accumulation of T1 CD8+ T cells, whereas the anti-PD-1+LAG-3 combination was associated with T3 CD8+ T cells and the CD4+ T cell cluster T2. Analysis of differentially expressed markers in the CD8+ T cell subsets identified T1 to express higher levels of cytotoxicity markers such as IFNγ, GZMB and PRF1 as well as multiple checkpoints including PDCD1 (PD-1), LAG-3, TIM3, TIGIT and CTLA-4 (**Figure 2D, E**). By contrast, the T3 subset, associated with anti-PD-1+LAG-3 responding tumors, had lower levels of cytotoxicity markers, decreased expression levels of LAG-3 and TIM3 (HAVCR2) and increased expression of TNFSF8, CXCR4 and IL21R (**Figure 2E**). IHC analysis confirmed the infiltration of CD4+ and CD8+ T cells (**Figure 2F**). We next utilized Chao1 analysis, to estimate the T cell receptor (TCR) diversity in the scRNA-seq data (25). We noted an expansion of TCR clones in tumors responding to PD-1 and CTLA-4, with the largest expansion seen in flank tumors responding to the PD-1+LAG-3 combination (**Figure 2G**). It thus appeared that anti-PD-1+LAG-3 drove the expansion of a limited number of T cell clones compared to the other ICI regimens.

### The anti-PD-1+LAG-3 and anti-PD-1+CTLA-4 combinations differentially polarize CD4+ T cells

Of particular interest were the CD4+ T cells, which comprised both T helper subsets and regulatory T cells (Tregs). Clustering analysis of the initial T2 cluster identified 2 further major subsets of CD4+ T cells (and a series of very minor clusters) in addition to Tregs (**Figure 3A**). The major CD4+ T cell cluster found in anti-PD-1+CTLA-4 responding tumors expressed markers consistent with Tregs such as FOXP3 and IL-2RA (CD25). It additionally expressed multiple checkpoints, including those not expressed in the other CD4+ T cell subsets, such as LAG-3. The T6 CD4+ T cell cluster were primarily found in anti-PD-1+LAG-3 treated tumors and expressed the highest levels of CXCR4, TNFSF8 and IL21R. This cluster also expressed genes known to be characteristic of Th1 cells (STAT1, STAT4, T-bet, IFN*γ*), Th2 (GATA3, STAT5, STAT6, IL4RA), Th9 cells (TGFRB2, IRF4, IL4Ra) and Th17 cells (IL21R, BATF, RORa, TGFBR2, STAT3) (**Supplemental Figure 2**). These genes had a heterogeneous expression, with some being found only in limited numbers of cells - suggesting that multiple phenotypes of helper CD4+ T cells were located within this cluster. Cytokine expression, such as IL17A, IL9, IL4, among others was poorly represented in the scRNA-Seq data, as others have previously reported (26). The final cluster T7, was comprised of a subset CD4+ T cells predicted to be activated that expressed CCR7, IL7R and TCF7 (**Figure 3A,B**). Flow cytometry demonstrated the anti-PD-1+LAG-3 combination to be associated with increased levels of CD69+ effector CD4+ T cells and reduced numbers of CD62L-CD4+ memory cells (**Figure 3C,D**). The increased levels of Tregs in the anti-PD-1+CTLA-4 treated SM1 flank tumors was confirmed both by flow cytometry and IHC staining for FOXP3 (**Figures 3C** and **3E**).

**Figure 3:**
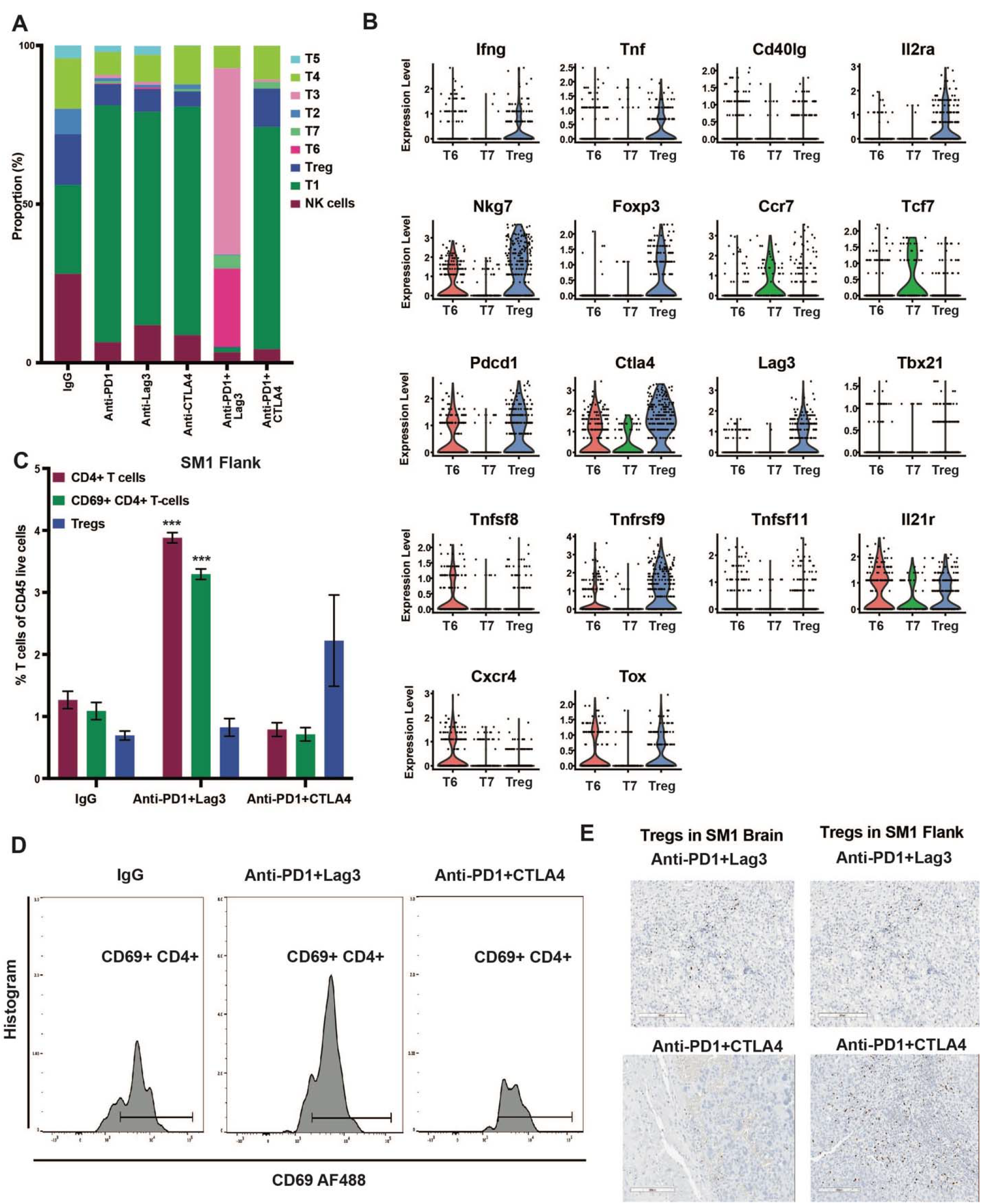
Anti-PD1+LAG-3 and Anti-PD1+CTLA-4 differentially polarize CD4+ T cells in the SM1 mouse melanoma model. A) Proportion of T cell clusters and three major subsets of CD4+ T-cells identified in the responding flank tumors from the indicated treatment groups. B) Expression of immune checkpoints and markers for T helper subsets and T regulatory cells in the three major subsets of CD4+ T-cells identified by violin plots. C) Percentage of CD4+ T cells and activated CD69+ CD4+ T cells in the responding flank tumors treated with combination of Anti-PD1+LAG-3 and Anti-PD1+CTLA-4 identified by flow cytometry. D) Flow cytometry histogram plot showing activated CD4+ T cells in tumors treated with IgG, Anti-PD1+LAG-3 and Anti-PD1+CTLA-4 as evidenced by cell surface CD69 and CD4 staining. E) Treatment with Anti-PD1+CTLA-4 increased Tregs infiltration in the responding flank tumors and relapsing brain tumors. The samples were stained with anti-FOXP3 by IHC. The results were represented as average ± SEM of 3 mice per group for panels C and D. Statistical significance was assessed with one-way ANOVA test (*, 0.001 ≤ p ≤ 0.05, **, 0.0001 ≤ p ≤ 0.001 and ***, p ≤ 0.0001).

### Anti-tumor responses to anti-PD-1+LAG-3 and anti-PD-1+CTLA-4 therapy show differential requirements for CD4+ T cells

As our scRNA-seq analyses suggested the anti-PD-1+CTLA-4 and anti-PD-1+LAG-3 combinations had differential effects upon T cell polarization we next determined the dependency of each ICI combination upon CD4+ and CD8+ T cell activity. It was noted that whereas depletion of CD8+ T cells abrogated responses to both anti-PD-1+LAG-3 and anti-PD-1+CTLA-4, CD4+ T cell depletion only impacted responses to the anti-PD-1+LAG-3 combination (**Figure 4A,B: Supplemental Figure 3**). These effects were noted in both the brain and flank tumors, with pronounced effects of CD4+ T cell depletion being noted in the brain metastases (**Figure 4A,B**). Similar findings were noted in the B16 melanoma model, with depletion of CD4+ T cells impacting responses to the anti-PD-1+LAG-3 combination, but not the anti-PD-1+CTLA-4 combination (**Figure 4C**).

**Figure 4:**
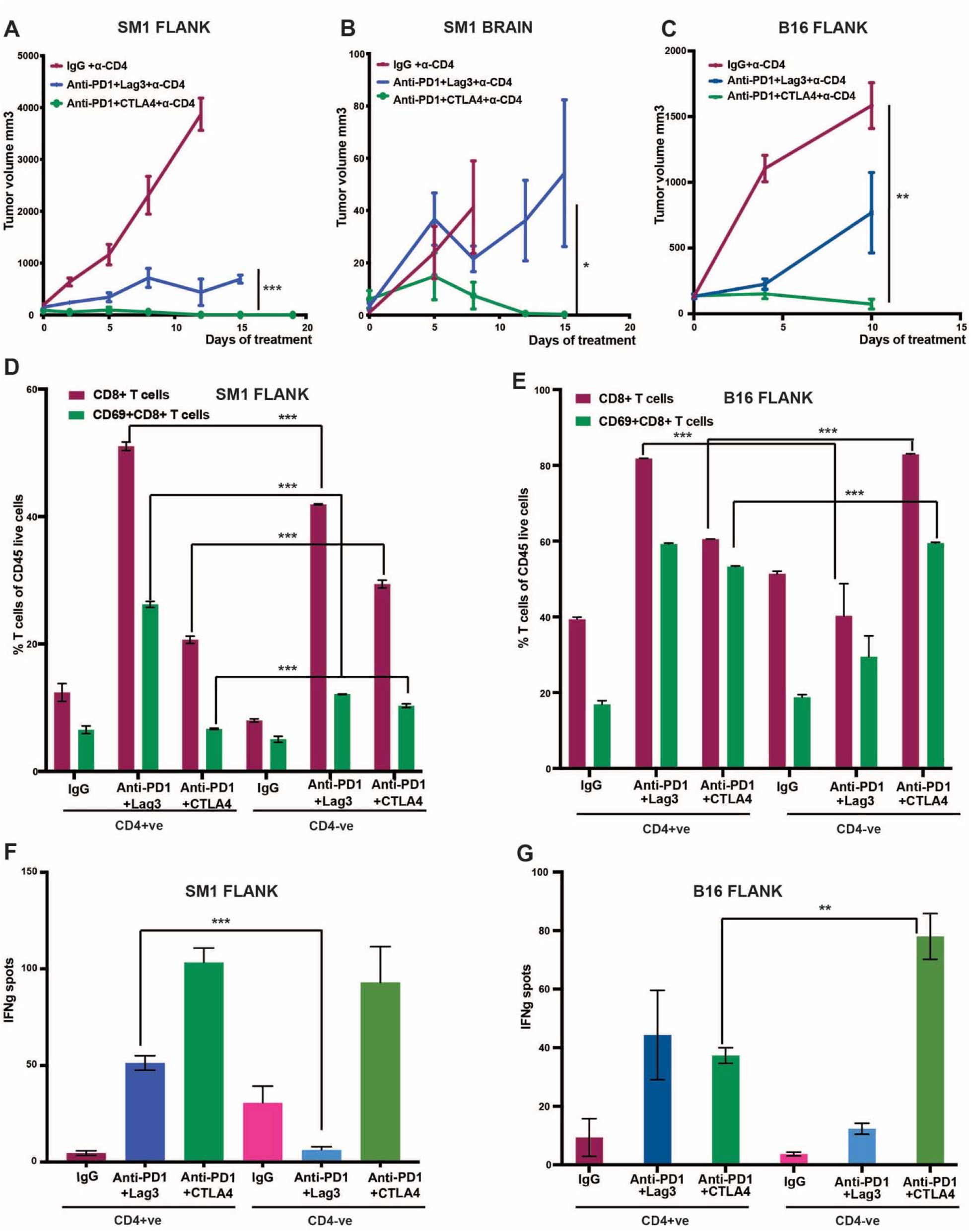
Responses to Anti-PD1+LAG-3 is dependent on CD4+ T cells. A, B) Depletion of CD4+ T cells demonstrates that responses to anti-PD1+LAG-3 but not PD1+CTLA-4 are dependent on CD4+ T cells in both SM1 flank and brain tumor models. The results were represented as an average ± SEM of 5 mice per group. C) Anti-tumor responses of the PD1+LAG-3 but not the PD1+CTLA-4 combination are dependent on CD4+ T cells in B16 flank tumors. Results show the average ± SEM of 5 mice per group. D, E) Decreased percentage of tumor-infiltrating CD69+CD8+ T-cells and CD8+ T-cells in SM1 and B16 flank tumors, respectively, following CD4+ T-cell depletion and anti-PD1+LAG-3 treatment. Increased percentages of CD69+CD8+ T-cells and total CD8+ T-cells were seen in SM1 and B16 flank tumors following CD4+ T-cell depletion and anti-PD1+CTLA-4 treatment. F, G) ELISPOT assays showing decreased levels of IFNg production following CD4+ T-cell depletion and treatment with the PD1+LAG-3 combination in both SM1 and B16 models. The results were represented as average ± SEM of 3 mice per group for panels D-G. Statistical significance was assessed with one-way ANOVA test (*, 0.001 ≤ p ≤ 0.05, **, 0.0001 ≤ p ≤ 0.001 and ***, p ≤ 0.0001).

There is evidence that CD4+ T cells exert anti-tumor effects through helper activity that increases CD8+ T cell function. Depletion of CD4+ T cells reduced the accumulation of activated CD69+ CD8+ T cells in anti-PD-1+LAG-3 treated tumors, suggesting a role for CD4+ T cell helper function in CD8+ T cell activation (**Figure 4D**). Conversely, depletion of CD4+ T cells in anti-PD-1+CTLA-4 treated tumors led to an increase in CD69+ CD8+ T cells, possibly a result of decreased Treg numbers in the anti-PD-1+CTLA-4 treated tumors (**Figure 4D: Supplemental Figure 4**). Similar findings were also noted in the B16 model, with depletion of CD4+ T cells being associated with decreased numbers of activated CD8+ T cells in anti-PD-1+LAG-3 treated tumors and increased CD8+ T cell activity in anti-PD-1+CTLA-4 treated tumors (**Figure 4E**). We next performed IFNγ ELISPOT assays on tumor infiltrating immune cells from the IgG and ICI treated SM1 tumors and demonstrated that the anti-PD-1+CTLA-4 combination led to greater increases in IFNγ release compared to the anti-PD-1+LAG-3 combination (**Figure 4F**). These findings confirmed the scRNA-seq data from Figure 2E suggesting that the T1 CD8+ T cells associated with PD-1+CTLA-4 treatment exhibited enhanced effector functions relative to the T3 subset of CD8+ T cells enriched for by the PD-1+LAG-3 combination. Consistent with the PD-1+LAG-3 combination being reliant upon CD4+ T cell help, it was noted that depletion of the CD4+ cells led to decreased IFNγ release from immune cells infiltrating anti-PD-1+LAG-3 treated SM1 and B16 tumors (**Figure 4F,G**), suggesting a role for T helper function. In contrast, depletion of CD4+ T cells from B16 tumor bearing mice treated with anti-PD-1+CTLA-4 led to an increase in INF*γ* release, suggesting a suppression of Treg activity (**Figure 4G**).

### Anti-PD-1+LAG-3 combination resistance in the MBM is associated with decreased T cell infiltrate and accumulation of MDSCs

Despite impressive responses at extracranial sites, similar numbers of mice failed anti-PD-1+LAG-3 and anti-PD-1+CTLA-4 therapy in the SM1 brain metastasis model. It was found that resistant tumors had a reduced proportion of CD8+ T cells compared to responding MBMs (**Figure 5A**). Levels of CD4+ T cells or Tregs did not significantly change in the resistant tumors. IHC analysis confirmed that the resistant tumors still contained CD4+ and CD8+ T cells (**Figure 5B**). We next performed tetramer assays to measure numbers of gp100+ (an antigen expressed by the SM1 cells) tumor-reactive T cells in tumors that were sensitive and resistant to anti-PD-1+CTLA-4 and anti-PD-1+LAG-3. It was noted that resistance to both ICI combinations was associated with loss of tumor reactive CD8+ T cells (**Figure 5C**). As suppressive myeloid cells can eradicate T cells in the tumor microenvironment (TME) we returned to our scRNA-Seq data to determine how tumor progression despite ICI combination therapy altered the repertoire of myeloid cells. Marked differences were seen in the myeloid composition of the resistant tumors treated with the single agent and combination ICI therapy (**Figure 5D**). It was found that the anti-PD-1+LAG-3 combination failures in the brain were highly enriched for MDSC-2, a cluster of putative myeloid-derived suppressor cells that expressed high levels of Arg1, NOS2, IRF7, HMOX1, CCRL2 (**Figure 5E** and **Supplemental Figure 5**). Flow cytometry analysis confirmed these findings and demonstrated an increased accumulation of Ly6C+ CD45+ M-MDSCS in mouse brain tumors that had relapsed on the anti-PD-1+LAG-3 combination (**Figure 5E** and **Supplemental Figure 6**). Other notable findings included the emergence of a cluster of macrophages-2 in the anti-PD-1 failures, and increased accumulation of the macrophages-1 and 2 in the anti-PD-1+CTLA-4 failures (**Figure 5D**). We next performed a cell-cell interaction analysis using SingleCellSignalR to infer the interactions between the MDSC-like cells in each sample and the identified T cell phenotypes. It was noted that the MDSCs in the anti-PD-1+LAG-3 combination failures had inhibitory interactions with the majority of the T cell phenotypes, suggesting that these cells may be responsible for the suppression of tumor-reactive T cell responses in the anti-PD-1+LAG-3 failure specimens (**Supplemental Figure 7**). As these interactions seemed to be bi-directional we depleted the CD4+ T cells from the IgG, anti-PD-1+CTLA-4 and anti-PD-1+LAG-3 treated mice and noted that suppressing CD4+ T cell accumulation was associated with increased MDSC numbers in the anti-PD-1+LAG-3 treated mice only (**Figure 5F**). Circos plots were utilized to illustrate bi-directional interactions between the MDSCs and T cells, demonstrating the high number of interactions in immune cells from anti-PD-1+LAG-3 treated tumors (**Figure 5G**). A proposed mechanism of action for the two different ICI combinations is shown in **Figure 6**.

**Figure 5:**
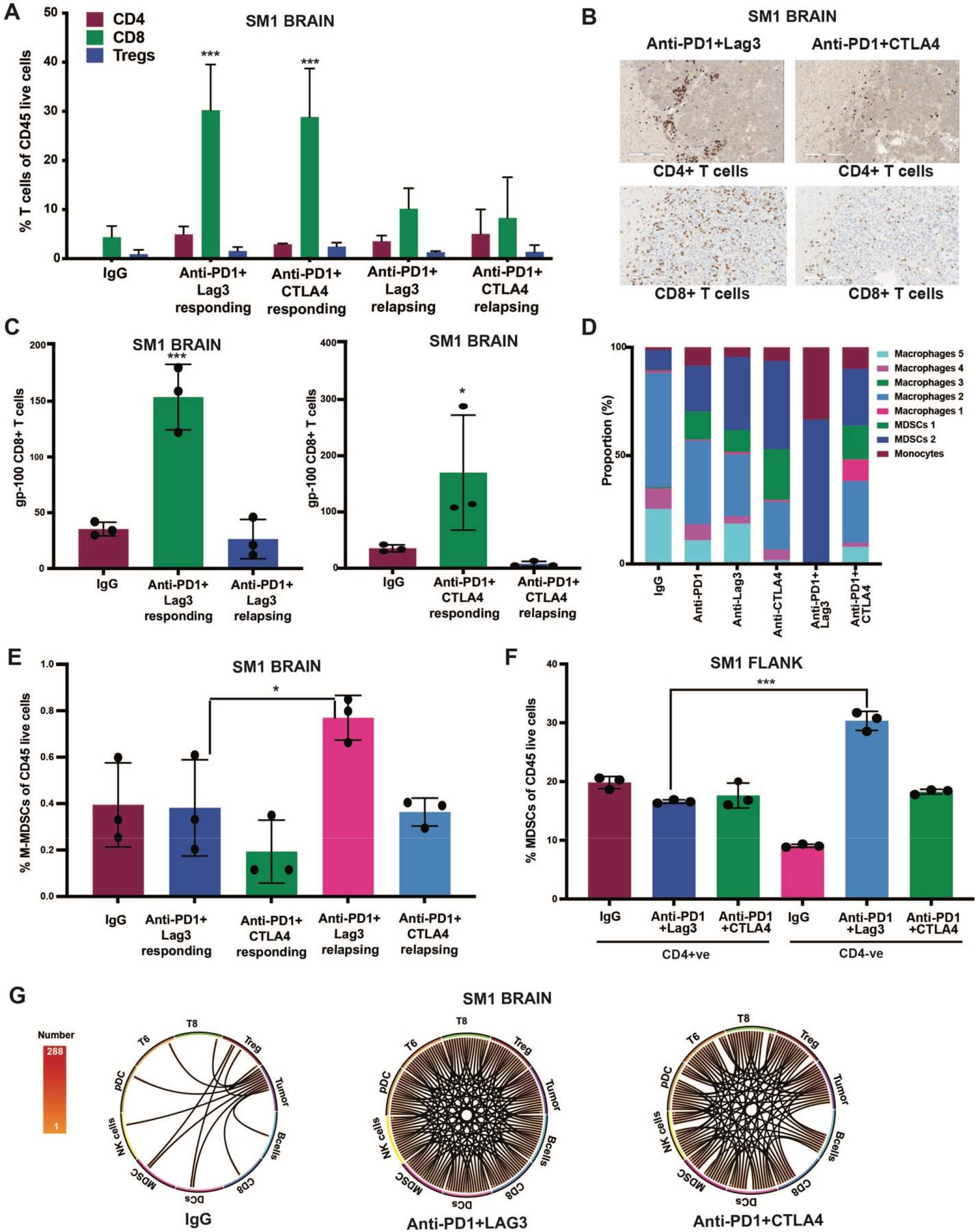
Escape from the PD-1+LAG-3 combination in the brain is associated with increased accumulation of MDSCs. A) Percentage of CD4+, CD8+ T-cells and Tregs in SM1 brain tumors following treatment with the PD1+LAG-3 and PD1+CTLA-4 combination. Decreased T cells infiltrate in resistant MBMs as compared to responding tumors. B) IHC analysis showing that resistant MBM still have some infiltrating CD4+ and CD8+ T-cells. C) The acquisition of resistance to each ICI combination is associated with loss of gp100-reactive T cells. Tetramer assays showing high levels of tumor reactive CD8+ T cells in responding MBMs, which then decline in resistant samples. D) scRNA-seq analysis of the myeloid cell proportions in MBM resistant to single agent and combination ICI shows enrichment for MDSCs in the PD-1+LAG-3 resistant brain tumors. E) Flow cytometry analysis shows increased tumor infiltrating MDSC numbers in MBM with PD-1+LAG-3 resistance vs. responding tumors. F) Depletion of CD4+ T cells is associated with increased MDSC numbers in SM1 flank tumors. Data shows flow cytometry analysis of MDSCs as a % of total CD45+ cells. G) Circos plots of directional cell-cell interaction analyses of the scRNA-seq shows increased interactions between MDSCs and T cells in the MBMs relapsing on PD-1+LAG-3 therapy. The results were represented as average ± SEM of 3 mice per group for panels A-F. Statistical significance was assessed with one-way ANOVA test (*, 0.001 ≤ p ≤ 0.05, **, 0.0001 ≤ p ≤ 0.001 and ***, p ≤ 0.0001).

**Figure 6:**
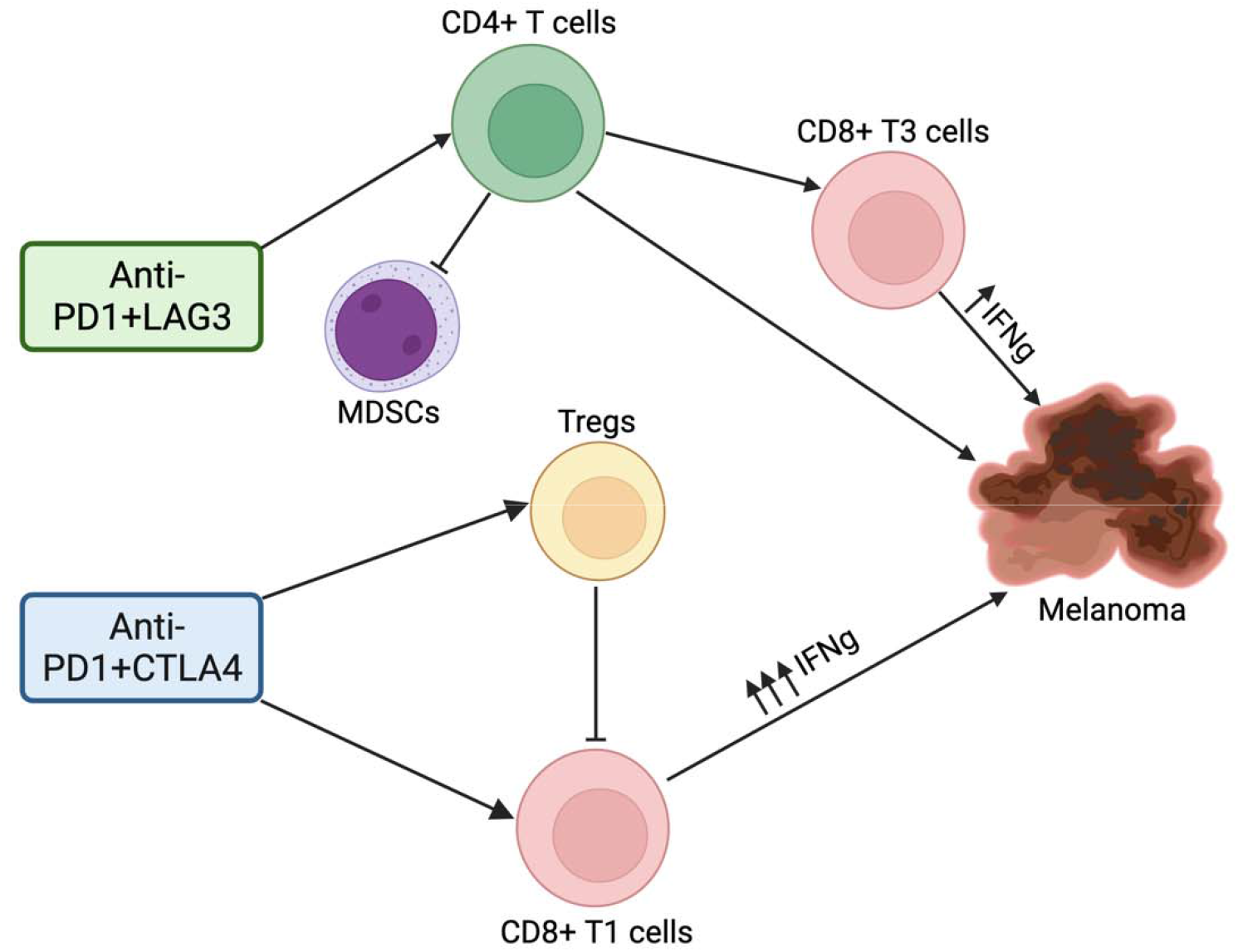
Proposed mechanism of the two ICI combinations.

## Discussion

CTLA-4, LAG-3 and PD-1 are all known to be expressed upon T cell activation and serve as biomarkers of T cell exhaustion. Despite these similarities, the mechanisms of action of the PD-1 +CTLA-4 and the PD-1+LAG-3 combinations have never been directly compared. In the current study we used two mouse melanoma models and high dimensional scRNA-seq analysis to identify a unique dependency of the anti-PD-1+LAG-3 combination upon CD4+ T helper cell function that was not seen in mice treated with the anti-PD-1+CTLA-4 combination. It was additionally found that the anti-PD-1+LAG-3 combination enriched for T cell transcriptional states that were very different from those associated with anti-PD-1 or anti-LAG-3 alone. Although the reactivation of CD8+ T cell activity has been the major focus of ICI research there is growing evidence that CD4+ T cells may also be critical (27). CD4+ T cells can differentiate into multiple T helper (Th) states including Th1, Th2, Th9, Th17 and Tregs that subserve different functions (28, 29). Th1 CD4+ T cells with cytotoxic function can mediate direct regression of established tumors in a major histocompatibility complex class II (MHC II)– restricted manner (28, 30, 31, 32). Cytotoxic CD4+ T cells have been identified in melanoma patients undergoing ICI treatment (33), and there is evidence that increased levels of circulating CD4+ T cells correlate with better outcomes in melanoma patients treated with anti-PD-1 therapy (34, 35). They also have important helper function that can improve B cell responses and CD8+ T cell responses directly (leading to memory) and indirectly through DC function (28, 29). Recent studies have also suggested roles for Th9 and Th17 T helper cells in anti-tumor responses. Th9 cells have been reported to have superior anti-tumor activity compared to other T helper subsets, and have the ability to activate both innate (NK cells, dendritic cells) and cytotoxic CD8+ T cell activity (34). Th17 cells also have anti-cancer activity, with studies showing that adoptively transferred Th17 cells can eradicate melanomas in mice (36, 37). There is also evidence that IL-17 deficient mice are more susceptible to melanoma development in the lung (37). Differentiation of Th17 cells requires TGF-β, IL-6 and IL-1beta and then IL-21 and IL-23 for their maintenance (38).

An analysis of the tumor infiltrating CD4+ T cell phenotypes identified key differences between the two ICI combinations. scRNA-Seq analysis demonstrated that CD4+ T cells from anti-PD-1+CTLA-4 treated tumors expressed multiple Treg markers including FOXP3 and IL2RA, whereas those from anti-PD-1+LAG-3 treated tumors were more characteristic of T helper CD4+ T cells. These T helper cells did not have one predicted phenotype and comprised cells with gene expression profiles previously associated with Th1, Th9 and Th17 helper cells. CD4+ T cell depletion studies confirmed this likely T helper phenotype, demonstrating a diminished response to anti-PD-1+LAG-3 therapy in two independent melanoma models, associated with reduced CD8+ T cell activation. Further evidence for a role for T helper activity came from the scRNA-seq-based cell type curation data which identified increased numbers of B cells and dendritic cells in tumors treated with the anti-PD-1+LAG-3 combination. In contrast, depletion of CD4+ T cells followed by anti-PD-1+CTLA-4 treatment had little effect upon anti-tumor activity and was instead associated with increased CD8+ T cell activity. This apparent paradoxical effect of CD4+ T cell depletion upon CD8+ T cell activation levels is likely consequence of the higher levels of Tregs in the PD-1+CTLA-4 treated tumors. It is known that Tregs express high levels of CTLA-4, and that CTLA-4 inhibition leads to Treg expansion (39). These effects are mediated through disruption of a CTLA-4 dependent feedback loop that leads to a CD28-mediated expansion of tumor-associated Tregs, increased tumor tolerance and decreased cytotoxic CD8+ T cell activity (39). Tregs limit cytotoxic CD8+ T cell anti-tumor activity through multiple mechanisms including the modulation of antigen presentation (through downregulation of CD80 and CD86 on DCs), the release of inhibitory cytokines (such as TGF-β and IL-10) and through depletion of important T cell growth factors in the environment (such as IL-2) (40). (41). LAG-3 has also been implicated in the regulation of Treg function, with prior work demonstrating that LAG-3 blockade inhibits the suppressive activity of Tregs *in vitro* and *in vivo* models of pulmonary vasculitis (42). Other studies have suggested that high levels of LAG-3 expression are critical for the maintenance of Treg suppressive function in models of mucosal immunity (43). In addition to its expression on T cells, LAG-3 is also expressed on multiple other immune cell types including B cells and γδ-T cells and it is known that mice who are LAG-3 deficient have increased numbers of T cells, B cells, macrophages, granulocytes and dendritic cells (44). Our scRNA-seq analysis of the T cell repertoire in melanomas treated with either anti-PD-1+CTLA-4 or anti-PD-1+LAG-3 led to the accumulation of distinct populations of both CD8+ and CD4+ T cells. The anti-PD-1+CTLA-4 combination enriched for a T1 population of CD8+ T cells, which expressed increased levels of cytotoxic markers such as IFN, GZMB and PRF1, whereas anti-PD-1+LAG-3 treatment was more associated with greater numbers of T3 cluster CD8+ T cells which expressed lower levels of cytotoxic markers. ELIPSOT assays confirmed these findings and showed that anti-PD-1+CTLA-4 treatment led to an increased infiltration of tumor-infiltrating IFN*γ* secreting CD8+ T cells compared to mice receiving anti-PD-1+LAG-3 therapy. It thus seemed that the anti-PD-1+CTLA-4 combination directly improved CD8+ T cell function, boosting their cytotoxic activity whereas the anti-PD-1+LAG-3 combination was associated with decreased Treg activity and enhanced CD4+ helper-mediated CD8+ T cell activation. The anti-PD-1+LAG-3 activated CD8+ T cells appeared to be less cytotoxic than those from tumors treated with the anti-PD-1+CTLA-4 combination. Although relatively little is known about the mechanism of action of the anti-PD-1+LAG-3 combination analysis of samples from the recent neoadjuvant phase 2 clinical trial of anti-PD-1+LAG-3 treated melanoma patients identified increased numbers of memory CD4+ T cells and CD8+ T cells in responding lesions, with responding patients additionally showing a reduction in M2-like macrophages (11).

Melanoma cells have multiple mechanisms to escape immune recognition (45, 46). Tumor intrinsic β-catenin activation in melanoma cells results in decreased dendritic cell (DC) accumulation, thus suppressing T cell priming (47). Other mechanisms can also include downregulation of antigen presentation via loss of expression of MHC-I expression (e.g. B2M) and acquired mutations in interferon/cytokine signaling pathways such as JAK2 mutations (48, 49). At the same time, resistance can arise through the accumulation of suppressive immune cell populations. This can be mediated through diverse subsets of myeloid cells, including cells resembling monocytic MDSCs, tumor associated macrophages (TAMs) and groups of tolerogenic DCs (tDCs). MDSCs and other immunosuppressive myeloid subsets are known to infiltrate melanoma samples and their presence is associated with poor responses to immune checkpoint inhibitor therapy (50, 51, 52). M-MDSCs, TAMs, and tDCs mediate their immunosuppressive activities through multiple mechanisms including the release of nitric oxide (NO), activity of arginase (ARG) I and II, production of TGF-β and the upregulation of immune checkpoints (such as PD-L1), among others (53, 54). Together, these effects inhibit the anti-tumor activities of anti-tumor NK and T cells and stimulate the recruitment and function of regulatory T cells (Tregs) (50).

Interrogation of scRNA-seq data from MBMs from SM1 melanoma cells relapsing/failing to respond to single agent or combination ICI therapy identified increased MDSC accumulation as a key feature of resistance to the anti-PD-1+LAG-3 combination in the brain. The subset of MDSCs identified expressed multiple genes known to limit T cell activity, and it was noted that the failure on anti-PD-1+LAG-3 therapy was associated with loss of tumor reactive gp100+ CD8+ cells. Although this loss of gp100+ CD8+ T cells was also noted in PD-1+CTLA-4 failures, increased numbers of MDSCs were not observed, suggesting an alternative mechanism of therapeutic escape. An analysis of inferred cell-cell interactions between the MDSCs and the T cell subsets identified multiple interactions, which appeared to be bi-directional. One unexpected consequence of CD4+ T cell depletion in anti-PD-1+LAG-3 treated tumors was an expansion of MDSCs, which were not observed when CD4+ cells were depleted in the anti-PD-1+CTLA-4 treated mice. It is entirely possible that activated CD4+ T cells could have been restraining MDSC expansion, although it remains to be determined whether this represents a direct or indirect effect. Together, our data suggest that the anti-PD-1+LAG-3 and anti-PD-1+CTLA-4 combinations have non-overlapping mechanisms of action and resistance mechanisms. Of particular note, the anti-PD-1+LAG-3 combination polarized the CD4+ T cells in a manner that was quite distinct from that of single agent anti-PD-1 or anti-LAG-3, as well as the anti-PD-1+CTLA-4 combination. It remains to be determined if this is a consequence of dual checkpoint inhibition driving T cell differentiation in a different manner to single checkpoint inhibition or whether this results from the targeting of different populations of T cells.

The observation that these two different, FDA-approved ICI combinations have different mechanisms of action is highly significant. Many patients exhibit upfront or acquired resistance to standard-of-care anti-PD-1 or the anti-PD-1+CTLA-4 combination, suggesting there could still be responses to second line ICI therapy with a different mechanism of action. It is also likely that the choice of ICI could be personalized, with patients who have MHC class 2 positive melanomas responding more favorably to anti-PD-1+LAG-3 therapy. As more therapeutic options become available for patients with advanced melanoma and melanoma brain metastases, selecting the optimal treatment regimen will become more critical.

