## Supplemental Figures 1-7 for "Defining the mechanisms of action and resistance to the anti-PD-1+LAG-3 and anti-PD-1+CTLA-4 combinations in melanoma flank and brain models"

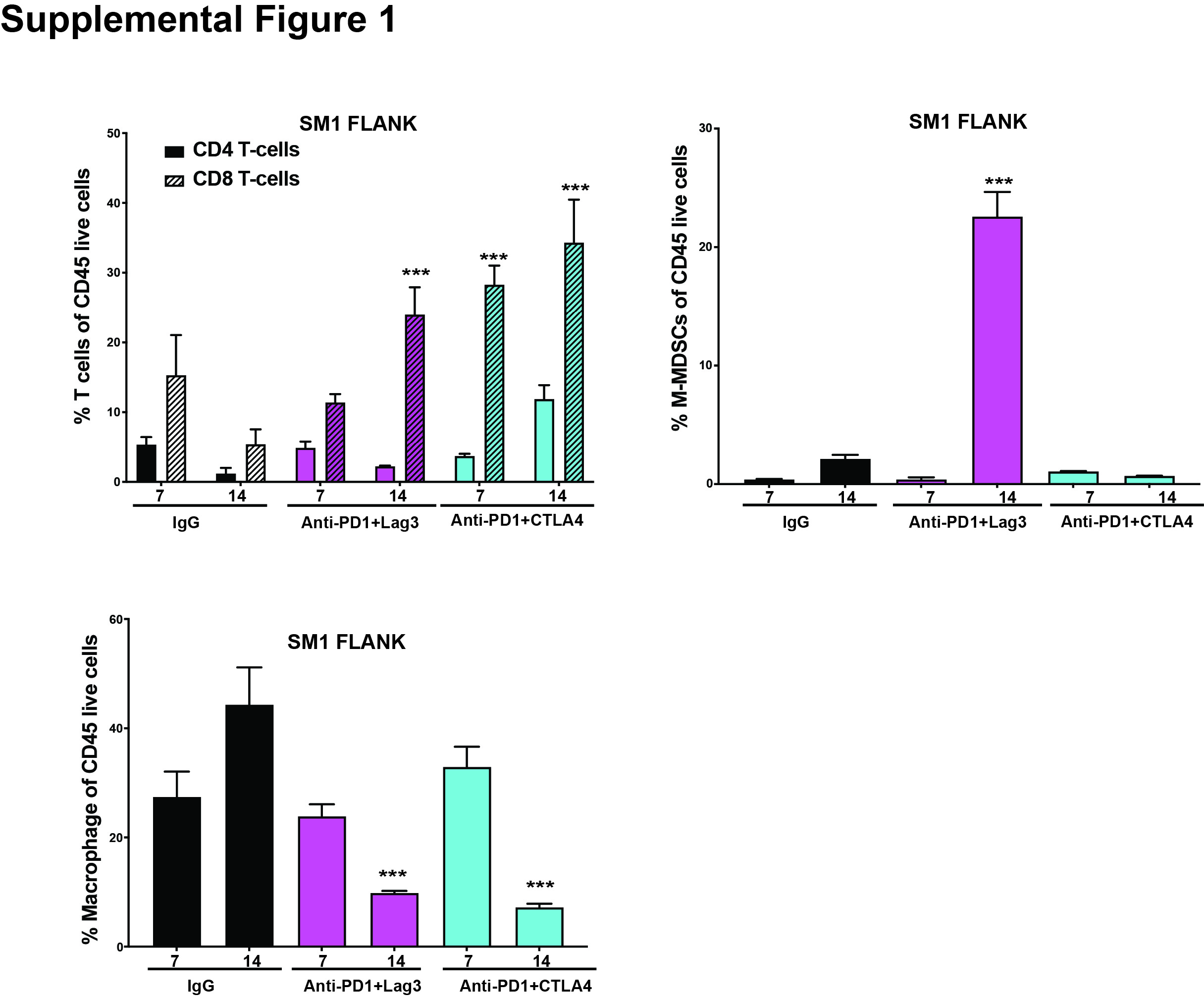

Kinetics experiment demonstrating increase in CD4+ and CD8+ T cells (left), increase in MDSCs on treatment with ICI combination therapy over a period of 14 days and a decrease in macrophages (bottom) on treatment over a period of 14 days in SM1 flank tumors. The results were represented as average ± SEM of 3 mice per group. Statistical significance was assessed with one-way ANOVA test (***, p ≤ 0.0001).

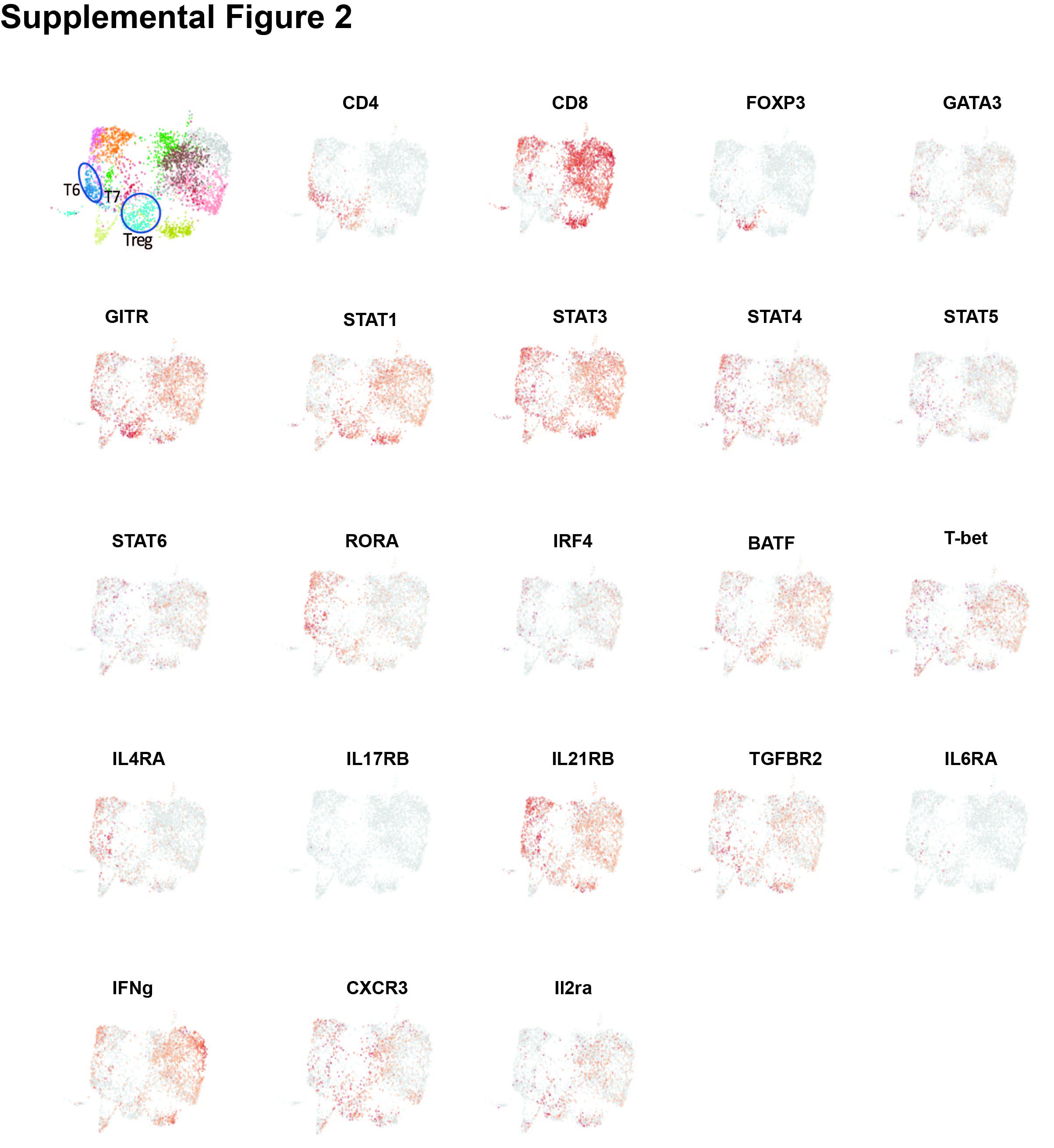

UMAPs illustrating increased expression of genes representing potential CD8+ T cells, Tregs and CD4+ Th1, Th2, Th9 and Th17 T cells in the T cell cluster.

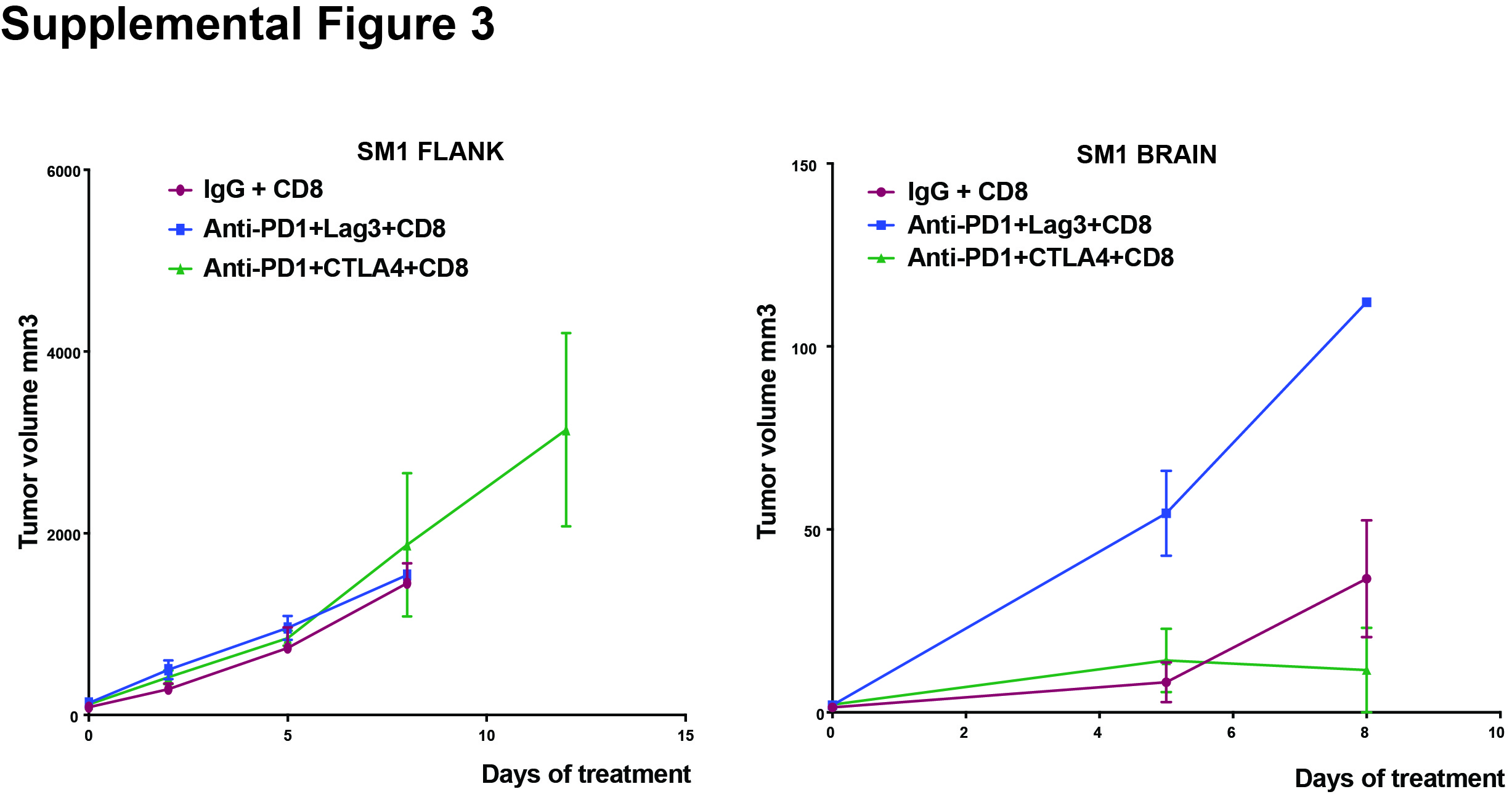

Responses to anti-PD1+LAG-3 and anti-PD1+CTLA-4 are dependent on CD8+ T cells in the flank and brain tumors, respectively, of the SM1 mouse melanoma model. Mice were treated with CD8-specific antibodies (100 μg/100 μL i.p) every 4 days before the tumor injections and before beginning the immunotherapy. The mice continued to receive Anti-CD8 along with IgG, anti-PD1+LAG-3 and anti-PD1+CTLA-4 during the entire course of treatment. The results were represented as an average ± SEM of 5 mice per group.

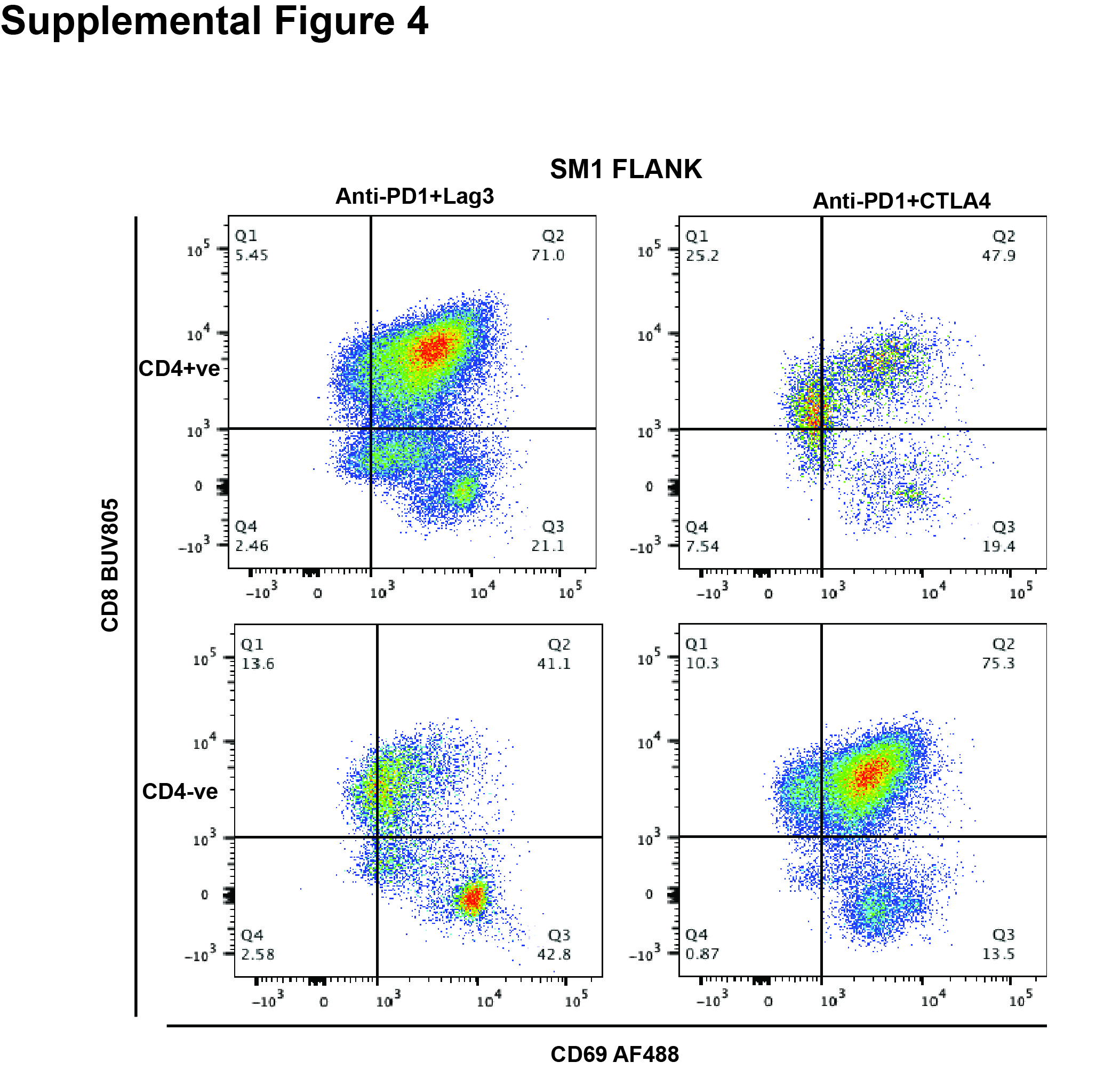

Flow cytometry pseudo color dot plot showing activated CD8+ T cells in tumors treated with anti-PD1+LAG-3 and anti-PD1+CTLA-4 following CD4+ T cell depletion as evidenced by cell surface CD69 and CD8 staining.

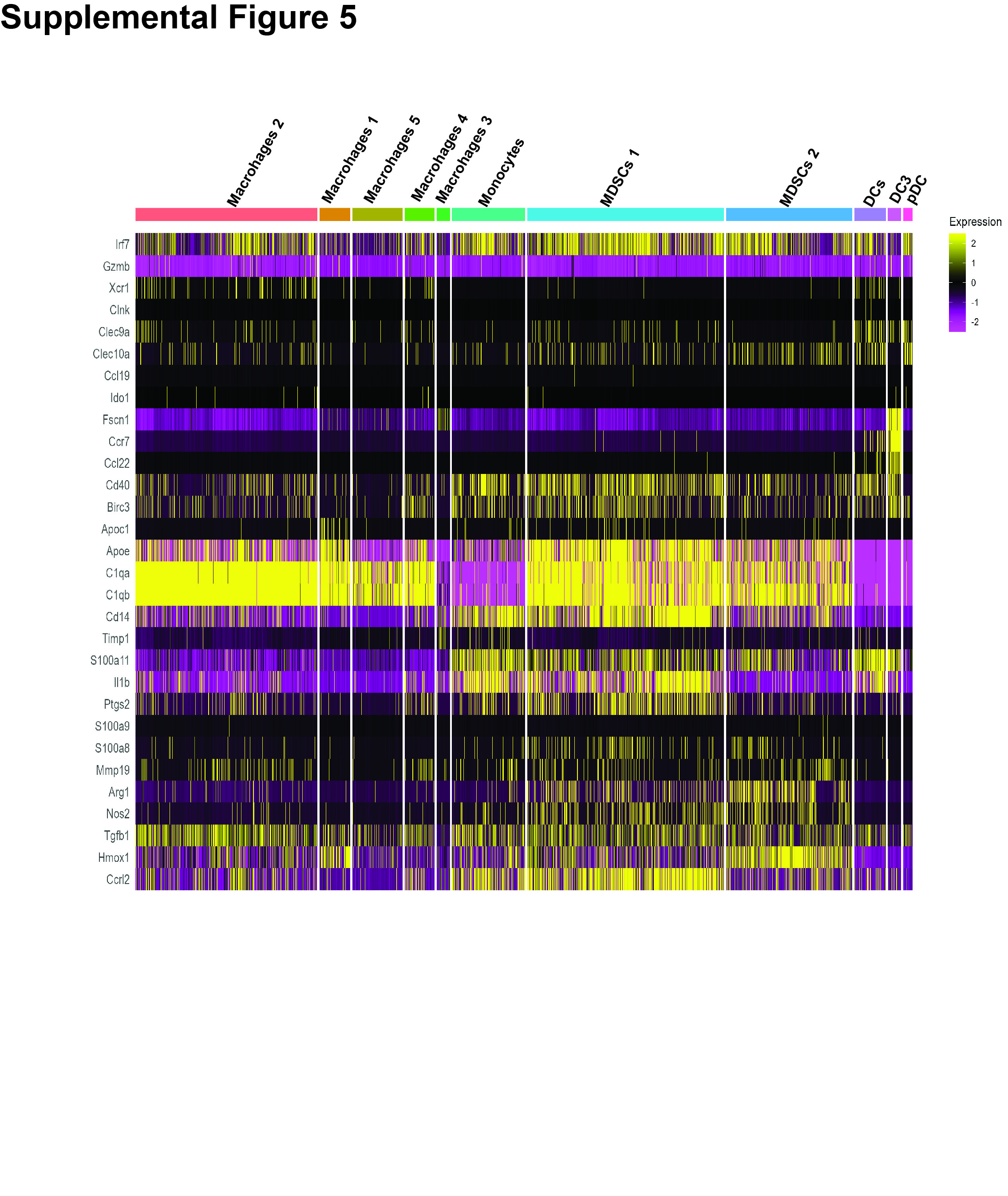

Expanded heat map with gene list showing unsupervised hierarchical clustering of macrophages, monocytes, MDSCs, DCs from scRNA-seq analysis of SM1 melanoma samples.

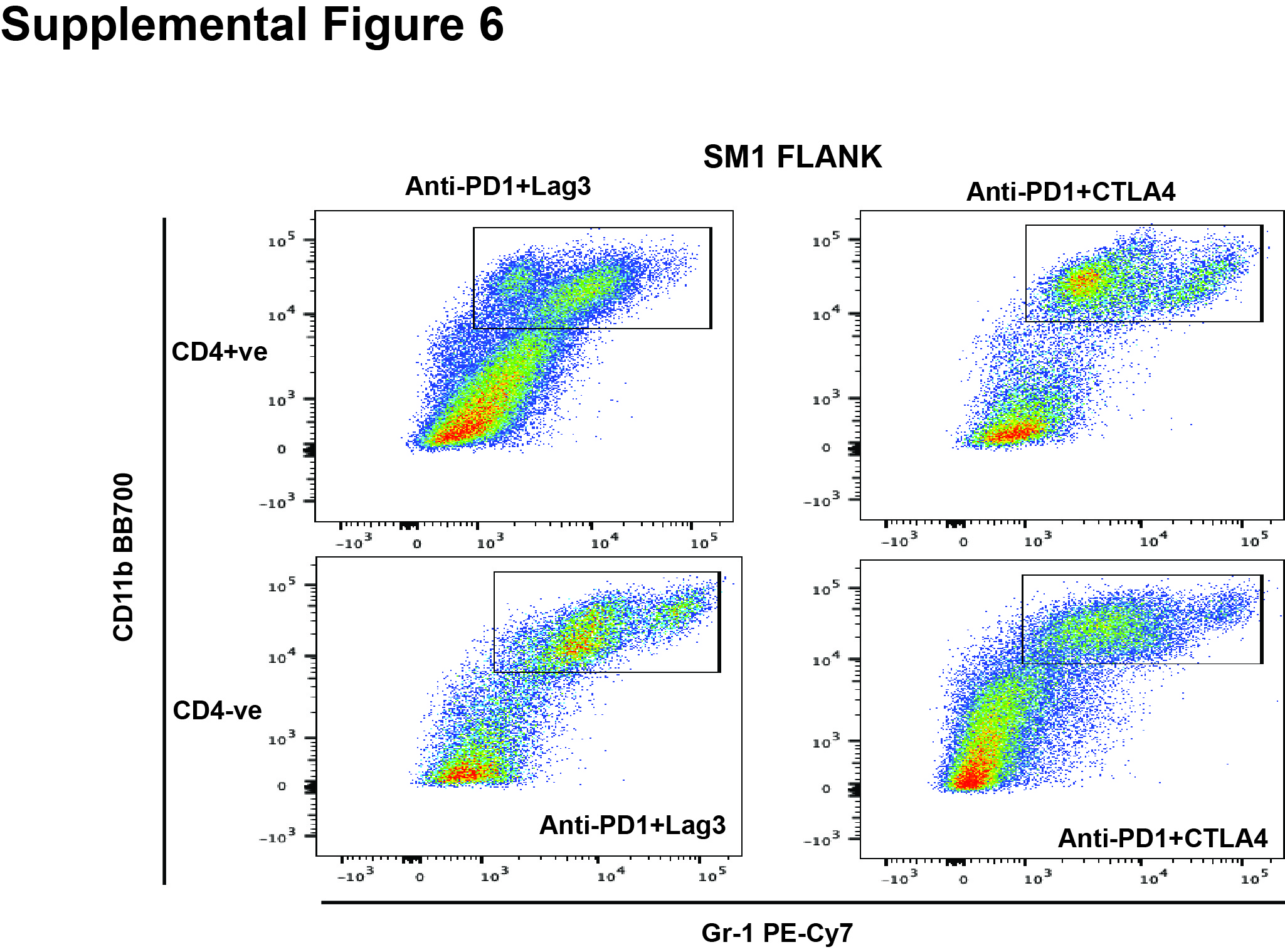

Flow cytometry pseudo color dot plot showing MDSCs in tumors treated with anti-PD1+LAG-3 and anti-PD1+CTLA-4 following CD4+ T cell depletion as evidenced by cell surface CD11b and Gr-1staining.

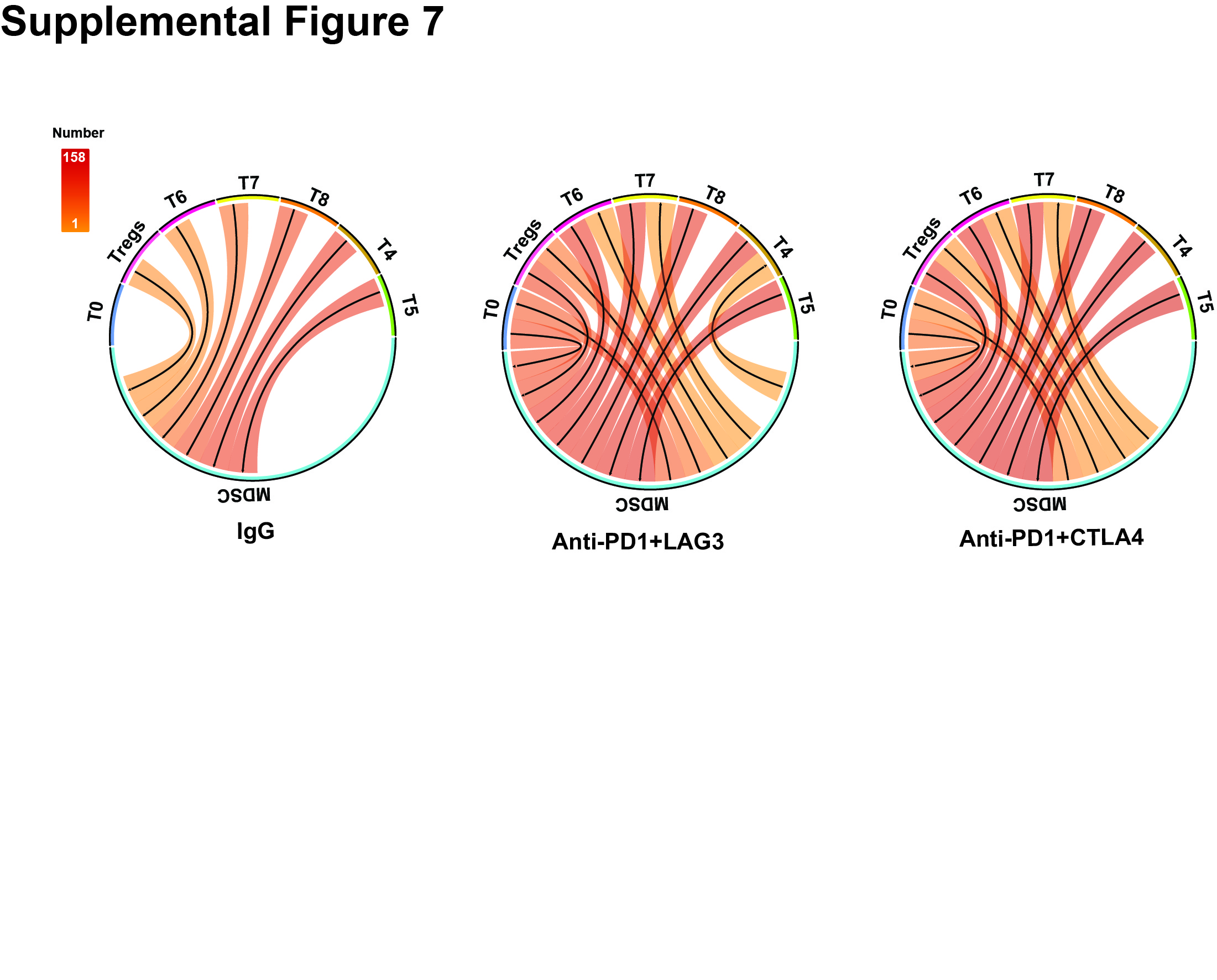

Cell-cell interaction analysis using SingleCellSignalR to deduce the interactions between MDSCs, and different clusters of T cells identified. Circos plots illustrating bi-directional interactions between immune cells and MDSCs across SM1 brain tumors resistant to anti-PD1+LAG-3 and anti-PD1+CTLA-4.
